# Wnt-3a exacerbates production of TNF-α in LPS stimulated microglia independent of the β-catenin canonical pathway

**DOI:** 10.1101/2025.06.30.662485

**Authors:** Gabrielle Federici, Sandy Stayte, Peggy Rentsch, Bryce Vissel

**Author notes:** Correspondence Bryce Vissel.

## Abstract

**Background:** Neuroinflammatory pathways are emerging therapeutic targets for neurological conditions such as Parkinson’s disease (PD). Studies have indicated Wnt-3a, a member of the wingless type MMTV integration (Wnt) signalling cascade, may exert anti-inflammatory effects via canonical pathway activation and β-catenin stabilisation. Furthermore, dysregulation of the Wnt/β-catenin pathway has been implicated in the degeneration of dopamine neurons in PD, however, stimulation of the canonical pathway via application of Wnt-3a to protect against inflammation and dopaminergic degeneration has not been explored.

**Methods:** Primary microglial cultures were stimulated with lipopolysaccharide (LPS) for 24 hours with or without Wnt-3a. TNF-α levels were measured via ELISA while changes in NFkB inflammatory pathway proteins and phosphorylated and non-phosphorylated β-catenin were analysed via capillary western blot. To assess Wnt pathway involvement, cultures were treated with DKK1 (β-catenin canonical pathway inhibitor), SP600125 (Wnt/Pcp pathway inhibitor) or U73122 (Wnt/Ca^2+^ pathway inhibitor). Finally, C57BL/6 mice received continuous intracerebroventricular infusion of Wnt-3a via osmotic pumps to investigate the effects of Wnt-3a on dopaminergic neuron survival and on microglial numbers in the MPTP model of PD.

**Results:** Wnt-3a alone had no effect on TNF-α release from microglia. However, when co-administered with LPS, there was a significant increase in cytokine release beyond that seen with LPS alone. Protein analysis revealed that this exacerbation in TNF-α levels was not due to alterations in the NFkB pathway or differences in activation of β-catenin. Furthermore, DKK1 treated cells showed no changes in TNF-α levels, however both SP600125 and U723122 were able to block Wnt-3a + LPS induced TNF-α release, implicating the non-canonical pathways. Meanwhile Wnt-3a *in vivo* did not alter dopaminergic or microglial populations in the substantia nigra in MPTP lesioned animals.

**Conclusion:** Together, these results suggest a pro-inflammatory response to Wnt-3a in an inflammatory context with little or no effect on resting microglia. Importantly, this outcome was independent of the β-catenin canonical pathway, revealing that Wnt-3a can increase pro-inflammatory TNF-α release via non-canonical signaling in an inflammatory environment. This demonstrates the importance of cellular context when identifying potential therapies for neurodegenerative diseases where neuroinflammation is a critical mediator of pathology.

## Introduction

The wingless type MMTV integration (Wnt) activated intracellular signalling cascade regulates multiple biological processes, ranging from embryogenesis and organogenesis to adult homeostasis. The Wnt family consists of 19 secreted Wnt ligands capable of activating three different pathways, often resulting in cell-type specific responses. The most extensively studied pathway within the Wnt signaling cascade is the canonical pathway, which relies on the activation of β-catenin. In the absence of Wnt ligands, β-catenin undergoes phosphorylation by the destruction complex, composed of GSK-3β and CK1, leading to its ubiquitination and subsequent degradation. However, when Wnt binds its receptor, it inhibits the destruction complex, causing β-catenin to stabilise and translocate into the nucleus to regulate the expression of multiple genes[1–3]. In contrast, the two non-canonical signalling pathways, the planar cell polarity pathway (Wnt/Pcp pathway) which is responsible for the remodelling of the cytoskeleton and orientation of cells, and the Wnt/Ca^2+^ pathway which releases intracellular Ca^2+^, operate independently of β-catenin [4, 5].

Dysregulation of Wnts has been observed in various diseases including epilepsy, cancer and metabolic diseases. Recent studies have suggested a connection between Wnt pathways and central nervous system (CNS) diseases, linking Wnt signalling to conditions such as schizophrenia, autism, mood disorders, Alzheimer disease (AD) and Parkinson’s disease (PD). Relevant to this study, Wnt, acting through β-catenin, has shown neuroprotective properties in a neuronal cell line treated with 6-OHDA, a neurotoxin mimicking Parkinsonian neurodegeneration[6], raising a question as to whether Wnts could also be neuroprotective *in vivo*.

Recently, studies have shown that Wnt activation can modulate the inflammatory response, including cytokine release, which is implicated in neurodegenerative disease pathology. In the brain, the classical innate immune response in microglia in the presence of pro-inflammatory stimuli such as LPS results from activation of toll-like receptors (TLRs) which in turn leads to activation of the IKKα/IKKβ complex within the cell and phosphorylation of IκB. Once phosphorylated, IκB is degraded, allowing NFκB to translocate into the nucleus where it activates various pro-inflammatory genes and contributes to the production of pro-inflammatory cytokines, including TNF-α.

Broadly, the Wnt canonical pathway is generally thought to trigger an anti-inflammatory response while the non-canonical pathways are thought to be pro-inflammatory [1, 2]. Moreover, the specific subgroup of Wnts appears to determine whether the canonical or non-canonical pathway is preferentially activated. The Wnt1 subgroup (Wnt2, Wnt3, Wnt3a and Wnt8) is thought to predominantly activate the canonical pathway, while the Wnt5a subgroup (Wnt4, Wnt5a, Wnt5b, Wnt6 Wnt7a and Wnt11) generally activates the non-canonical PCP or Ca^2+^ pathway[2]. However, this classification represents a simplification of the complex signalling interactions, since the same Wnt protein can activate different pathways depending on the subtype of receptor, the cell type involved and the specific cellular context. Nevertheless, the question follows as to whether Wnts that preferentially stimulate the β-catenin pathway, which is thought to be anti-inflammatory, holds potential for modulating inflammatory responses in disease.

Among the large family of Wnt ligands, Wnt-3a was the first to be purified [7, 8] and is perhaps the most extensively studied of the Wnts, and is among those that is thought to preferentially stimulate the canonical β-catenin pathway. The objective of this study was to investigate how inflammation is modulated by Wnt-3a in resting and activated microglia and to determine the molecular pathways involved. Furthermore, the effects of Wnt-3a were investigated in the MPTP mouse model of PD, a well-established model characterised by neuroinflammation, neurodegeneration and dysregulation of the Wnt pathway[9]. Our findings reveal an increase in neuroinflammation following application of Wnt-3a to reactive inflammatory cells, which is a result of signaling through non-canonical pathways.

## Materials and Methods

### Animals

Male C57BL/6 mice aged 11 weeks, and time mated SwissTacAusb female mice aged between 9-11 weeks and 14 days pregnant, were obtained from Australian BioResources (Moss Vale, Australia). Upon arrival, the mice were housed for a week to acclimate to the new environment before experiments began. The C57BL/6 mice were initially housed in groups of maximum 5 individuals per cage. Once the study started, they were individually housed to optimise the recovery process and ensure the integrity of the surgical sutures of each individual mouse. The mice were kept on a 12-hour light/dark cycle with access to food and water *ad libitum*. All animal procedures were performed with the approval of the Garvan Institute Animal Ethics Committee under approval numbers 18/16 and 17/35, in accordance with the Australian National Health and Medical Research Council animal experimentation guidelines and the local Code of Practice for the Care and Use of Animals for Scientific Purposes (2004).

### Primary microglia cell culture

Primary microglia were prepared as previously described with minor modification[10]. Briefly, the olfactory bulbs and the cerebellum of P0-P2 SwissTacAusb pups were removed and the cortices of each brain were dissected followed by the removal of the meningeal tissue. Once all the cortices were dissected, 2.5 mL of trypsin/EDTA (Thermo Fisher, Australia) was added and left for 30 minutes in a humidified incubator with 5% of CO_2_ at 37°C. A standard medium consisting of 10 mL of Dulbecco’s modified Eagle’s medium (DMEM)/F12 (Thermo Fisher scientific, Australia) supplemented with 10% foetal bovine serum (FBS, Sigma Aldrich, Australia) and 1% penicillin/streptomycin (Sigma Aldrich, Australia) was added to stop the reaction. Cells were then centrifuged at 1200 RPM for 5 minutes and the pellet resuspended in the standard medium to obtain a single cell suspension consisting of both microglia and astrocytes. The cells were then transferred into a T75 flask with preparation of 4 pups giving approximately 10-15 × 10^6^ glial cells per T75 flask. Finally, culture media was changed the day after and every 2–3 days thereafter. After 10–11 days, when cells had reached confluence, microglia were separated from the astrocytes by shaking the flask for 6 hours at 200 RPM. Following shaking, the media was collected and centrifuged at 1200 RPM for 5 minutes. The pellet, containing microglia, was resuspended in standard culture medium and plated at 3 × 10^6^ cells/well in a 12-well plate that had been pre-coated with Poly-D-lysine (PDL, Sigma Aldrich, Australia) 4 hours prior. Cultures were monitored daily under a microscope and experiments were started when the cells reached 70-80% confluence.

### Cell stimulation and collection

Microglia were left in a serum-free medium for 4 hours and then treated with 1 μg/mL LPS (Sigma, USA) and/or 300 ng/mL Wnt-3a (R&D Systems, USA) for 24 hours, with PBS acting as a control. The media was then collected and centrifuged at 12,000 RPM for 10 minutes at 4°C with supernatants collected and transferred into new tubes and stored at −80°C until use. The cells on the bottom of the plate were washed for 5 minutes with 1x PBS and then lysed for 30 minutes in RIPA buffer containing protease and phosphatase inhibitors (1:100, Sigma, Australia). After 30 minutes, cells were scraped and centrifuged at 12,000 RPM at 4°C for 10 minutes. Supernatants were transferred into new tubes and stored at −80°C for future experiments.

### Canonical/non-canonical pathway inhibition

To investigate the involvement of the Wnt pathway in inflammation, three inhibitors (or DMSO/PBS control) were added to the cultures 30 minutes prior to LPS and/or Wnt-3a stimulation. Mouse recombinant DKK1 (20 µg/mL, R&D systems) was added to inhibit the canonical pathway, while the JNK inhibitor SP600125 (15 µg/mL, Sigma Aldrich) was added to inhibit the PCP pathway. Additionally, the PLC inhibitor, U73122 (0.5 µg/mL, Sigma-Aldrich) was added to inhibit the Ca^2+^ pathway.

### Capillary western blotting (Wes)

Differences in protein expression were determined using an automated capillary western system (Wes, Protein simple, San Jose, CA). Using the Wes 12–230 kDa separation module (ProteinSimple, SM-W004) lysates of cells were diluted 1:100 in the 0.1x sample buffer at 0.25 mg/mL, prepared according to manufacturer’s instructions. The following primary antibodies were utilised: β-catenin (Abcam, cat # Ab 32572 at 1:10), non-phosphorylated β-catenin (Ser33/37/Thr41) (Cell signalling, cat # 4270s at 1:10), phosphorylated-GSK-3β (S9) (Cell signalling, cat # 9323s at 1:25), phosphorylated-GSK-3β (Tyr216) (Abcam, cat # Ab75745 at 1:10), NFκB p65 (Cell signalling, cat # 4764s at 1:20), phosphorylated-NFκB p65 (Ser 536) (Cell signalling, cat # 3033s at 1:20), phosphorylated-IKKα (Ser176)/IKKβ (Ser177) (Cell signalling, cat # 2078s at 1:10), phosphorylated IκB-α (ser32) (Cell signalling, cat # 2859s at 1:10), GAPDH (Cell signalling, cat # 2118L at 1:20). Secondary antibodies used were part of the anti-rabbit detection module (ProteinSimple, DM-001), the anti-mouse detection module (ProteinSimple, DM-002) or the total protein detection module (ProteinSimple, DM-TP01). Data were analysed using the compass software (compass for Simple Western, version 4.0.0) which displays the electropherogram and a virtual blot-like image. The area under the curve was used to calculate the ratio of protein/GAPDH, except for IκB-α which was compared to the total protein level as its molecular weight was similar to GAPDH.

### TNF-α ELISA

Sandwich enzyme-linked immunosorbent assays (ELISA) were performed on the culture medium of primary microglia following the manufacturer’s instructions (Elisa max deluxe set TNF-α Biolegend). Results were normalised to the total amount of protein measured via Bradford assay. All samples were run in duplicate.

### Osmotic pump preparation and implantation

For all in vivo experiments, osmotic micro-pumps (Model 1007D; Alzet, Palo Alto, California) were filled with a volume of approximatively 90-100 µl of Wnt-3a (R&D systems, 0.5 ng/µl in water + 0.1% BSA) or vehicle. C57BL/6 animals were anaesthetised with a mixture of ketamine (8.7 mg/ml, Mavlab, Slacks Creek, QLD) and xylazil (2 mg/ml, Troy Laboratories Pty Ltd, Smithfield, Australia) and placed in a mouse stereotaxic apparatus (Kopf Instruments). The micropumps were implanted subcutaneously along the back of the neck and connected to a cannula inserted into the ventricles at the following co-ordinates relative to bregma; anterior-posterior (AP) = −0.2mm, medial-lateral (ML) = +1.0 mm and dorsal-ventral (DV) −2.8 mm.

### MPTP administration

The day after osmotic pump implantation surgery, C57BL/6 animals received subcutaneous injections of 20 mg/kg of MPTP (Sigma Aldrich) or saline control every 2 hours for 8 hours (4 injections per animal), as described previously[11, 12]. All mice were euthanised 8 days after the final MPTP/saline administration.

### Immunohistochemistry

C57BL/6 mice were anaesthetised with a ketamine/xylazil mixture and transcardially perfused with ice cold 4% paraformaldehyde (PFA, Sigma Aldrich, Australia) dissolved in PBS. Brains were then dissected out and post-fixed overnight in 4% PFA at 4°C and then cryoprotected in 30% sucrose for 2-3 days at 4°C. Finally, tissues were frozen at −80°C in OCT compound (Tissue-Tek^®^; Sakura, Torrence, CA, USA) and cryosectioned coronally at 40 μm. Free floating sections were incubated in the following primary antibodies: monoclonal mouse tyrosine hydroxylase (TH, 1:1000 Sigma Aldrich cat # T2928), monoclonal mouse neuronal nuclei (NeuN 1:500, Merck Millipore, cat # MAB377) or polyclonal rabbit Iba1 (1:1000, Novachem, cat # 019–19741) for 72 hours at 4°C. The sections were then incubated in biotin-labelled secondary antibodies (1:250, anti-rabbit biotin, Thermo Fisher Scientific, Australia, cat # B2770; 1:250 anti-mouse biotin, Abcam, USA, cat # ab6788) overnight at 4°C followed by avidin-biotin complex (Vector Laboratories) incubation for 1 hour at room temperature. 3,3’-Diaminobenzidine (DAB, Abacus) was used to detect TH immunolabeling. NeuN and Iba1 immunolabeling was detected with DAB intensified with nickel ammonium sulphate and counterstained with polyclonal rabbit anti-TH (1:1000, Merck Millipore cat # AB152) or mouse monoclonal TH (1:1000) that was detected with Nova-Red (Vector Laboratories, USA) to outline the substantia nigra.

### Stereology

Quantification of neuronal and microglial cell populations was conducted using the optical fractionator method using Stereo Investigator 7 software (MBF Bioscience). The region of interest was traced using the 10X objective and the cells were counted using the 100X objective. The guard zone height was 5 μm and the optical dissector height was 10 μm. For the estimations of TH positive populations, the counting frame was 60 μm x 60 μm and the grid size was 113 μm x 73 μm. For the estimations of NeuN positive cells, the counting frame was 65 μm x 65 μm and the grid size was 155 μm and 125 μm. For the estimation of Iba1 positive populations, the counting frame was 58 μm x 58 μm and the grid size was 61 μm x 61 μm. The average thickness of the section was determined at a minimum of 5 sites. For TH and NeuN, every third section, to a total of 10 sections to encompass the entire SNpc were counted. For Iba1, every sixth section to a total of 5 sections were counted. The coefficient of error was calculated following Gundersen and Jensen and errors ≤0.10 were acceptable[13]. The SNpc was defined from −2.8 to −3.88 mm relative to bregma based on the Paxinos atlas for the mouse brain[14].

### Statistical analysis

All statistical analysis was performed with GraphPad Prism version 8.0.2 (GraphPad Software, Inc). Prior to assessing differences between the mean, the data was tested for normality using Shapiro-Wilk test. For normally distributed data, differences between means were assessed, as appropriate, by one-way or two-way ANOVA followed by Tukey or Sidak *post hoc* analysis. For non-normally distributed data, differences were assessed by Kruskal-Wallis test followed by Dunn’s *post hoc* analysis. All data is presented as mean ± standard error of the mean (SEM). For all data, significance was defined as *p < 0.05, **p < 0.01 and ***p < 0.001.

## Results

### Wnt-3a increases inflammation in reactive microglia primary cultures

The effects of Wnt-3a were analysed in both resting and inflammatory environments. To stimulate the inflammatory response in glial cells, we applied LPS (or saline control) with or without coadministration of Wnt-3a. Within glial cells, inflammatory stimuli leads to the IKKα/IKKβ complex becoming activated within the cell, leading to the phosphorylation of IκB, an inhibitor of NFκB. Once phosphorylated, IκB is degraded, allowing NFκB to translocate into the nucleus where it activates various pro-inflammatory genes and contributes to the production of pro-inflammatory cytokines, including TNF-α. As expected, LPS led to a significant release of TNF-α from microglia compared to unstimulated cells. Interestingly, while Wnt-3a alone did not increase the level of TNF-α compared to saline controls, when applied with LPS, Wnt-3a substantially exacerbated the release of TNF-α compared to cells stimulated with LPS alone (Figure 1A). When examining classical inflammatory pathways upstream of this process, our results revealed that the levels of active IKKα/IKKβ (Figure 1B) and IκB (Figure 1C) increased in response to LPS but were not further increased in LPS + Wnt-3a treated cells. Similarly, while the total level of NFκB remained unchanged across all groups (Figure 1D), the level of NFκBSer536, representing active NFκB, increased in response to LPS but was not further increased by the co-administration of Wnt-3a (Figure 1E). Surprisingly, Wnt-3a alone also increased the level of active NFκB compared to the control in this cell type (Figure 1E), suggesting some activation of this pathway has occurred, though this did not lead to increased TNF-α release. It is interesting to note that no differences were observed between the LPS and LPS + Wnt-3a group in any of these proteins, suggesting that Wnt3-a exacerbates release of TNF-α via a mechanism independent of increased activation of IKKα/IKKβ or phospho-NFκB.

**Figure 1:**
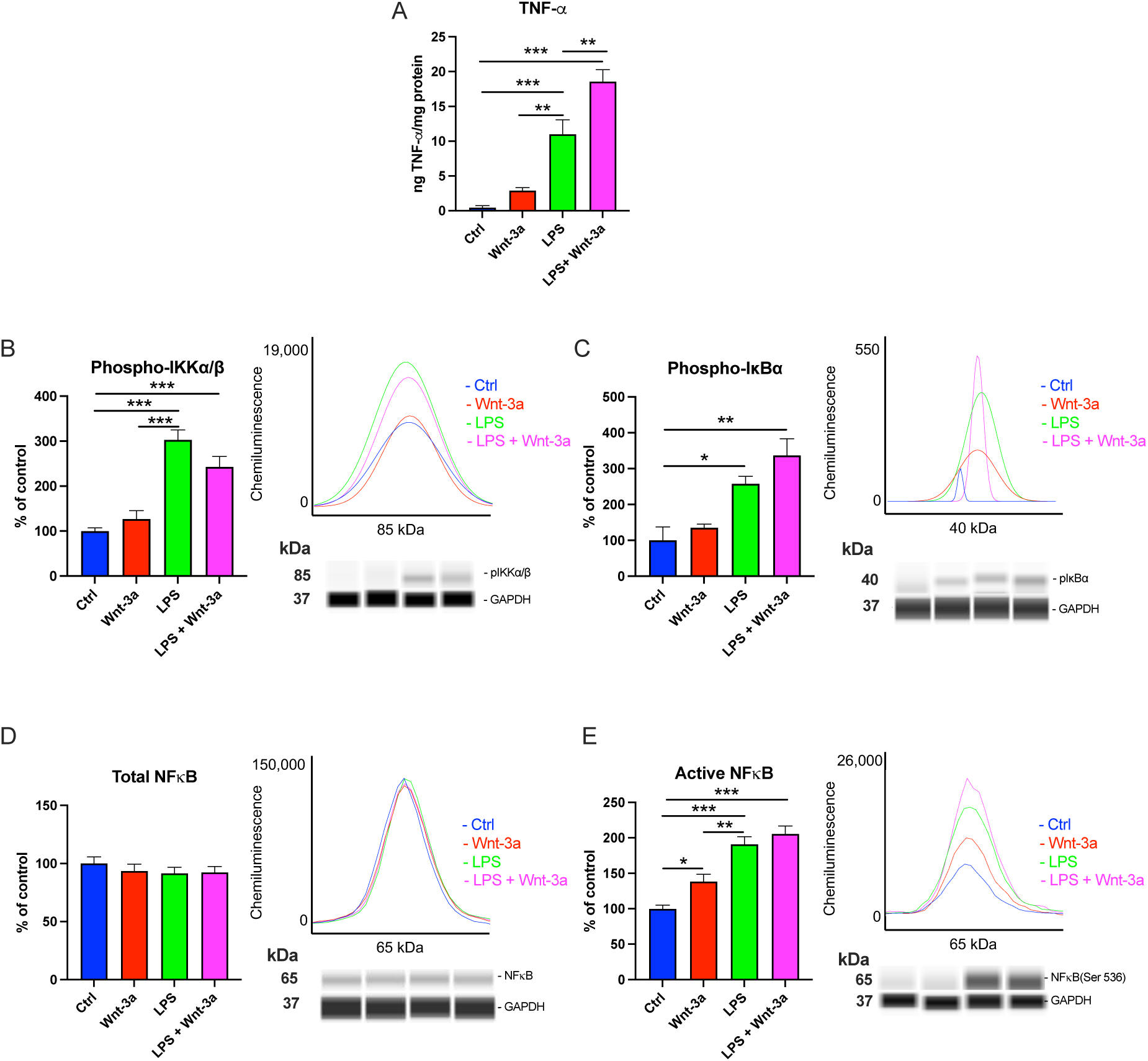
Changes in NFκB pathway proteins in primary microglia cultures stimulated with LPS and Wnt-3a. (A) Level of TNF-α produced after 24 hours of LPS and/or Wnt-3a stimulation N=10/group. Relative changes and corresponding Wes-generated electropherograms and pseudobands of (B) phospho-IKKα/β (C) phospho-IκBα (D) total NFκB and (E) active NFκB in microglia stimulated with LPS and/or Wnt-3a. All values are shown as mean ± standard error of the mean (SEM). *p<0.05, **p<0.01, ***p<0.001.

### Wnt-3a mediated exacerbation of inflammation is independent of the canonical pathway

Wnt-3a triggers the canonical pathway by binding to cell surface Frizzled (Fz) and LRP5/6 coreceptors, ultimately preventing β-catenin being targeted for degradation in the cytosol by the destruction complex via phosphorylation at Ser33, Ser37, and Thr41 sites[15]. This allows β-catenin to subsequently stabilise and translocate to the nucleus, activating Wnt target genes, some of which have an anti-inflammatory effect [16–18]. Therefore, to determine if the pro-inflammatory effects of Wnt-3a when administered in conjunction with LPS were due to alterations in the expression of phosphorylated vs non-phosphorylated β-catenin, we analysed both total β-catenin and active β-catenin, respectively, in our treatment groups.

While Wnt-3a alone did not significantly alter total β-catenin expression in microglia (Figure 2A), there was a significant reduction in β-catenin in LPS treated cells compared to Wnt-3a treated cells (p<0.05) suggesting some additional degradation of β-catenin in an inflammatory context, however it should be noted no statistical difference was found between LPS and control groups (p=0.0989). Interestingly, we found that LPS + Wnt-3a treated cells had significantly higher expression of total β-catenin compared to cells exposed to LPS alone, however we found no difference in β-catenin expression between this group and vehicle controls (p=0.3116), indicating these changes are unlikely to explain the exacerbation of TNF-α release in microglia following this co-stimulation.

**Figure 2:**
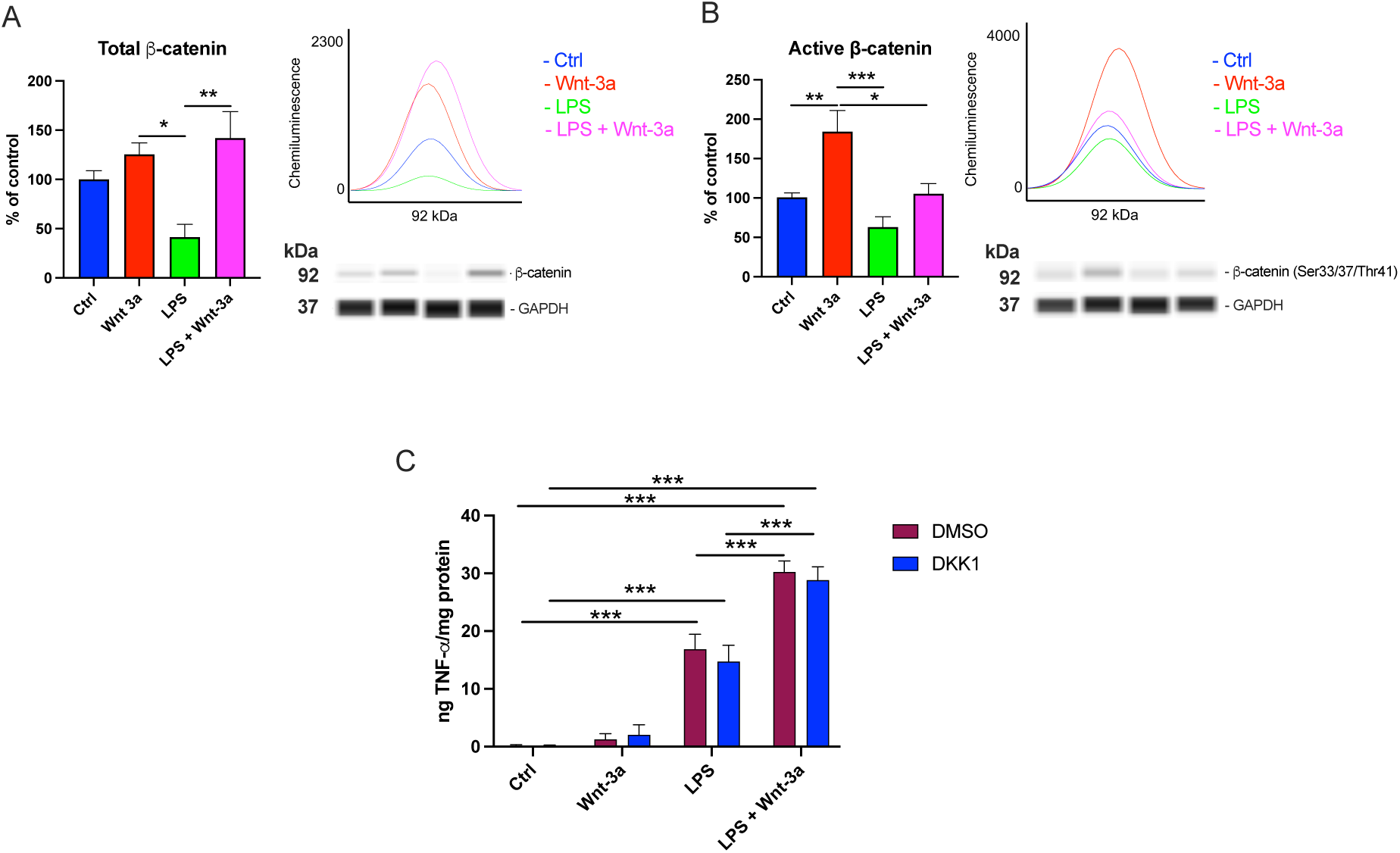
Canonical pathway response to Wnt-3a stimulation. Relative changes and corresponding electropherograms and Wes-generated pseudobands of (A) total β-catenin (B) active β-catenin in microglia stimulated with LPS and/or Wnt-3a (n=5-7/group). (C) Analysis of TNFα levels in microglia stimulated with DKK1 or PBS vehicle via ELISA (n=5/group). All values are shown as mean ± standard error of the mean (SEM). *p<0.05, **p<0.01, ***p<0.001.

As expected, there was a significantly increased level of active β-catenin, represented by increased expression of β-catenin(Ser/33/37/Thr41), in Wnt-3a treated microglia and thus confirming activation of the canonical pathway following Wnt-3a stimulation (Figure 2B). While no difference was found between cells stimulated with LPS alone or co-stimulated with LPS + Wnt-3a, active β-catenin levels were suppressed in both of these groups when compared to Wnt-3a stimulated cells. This suggests alterations of the canonical pathway in an inflammatory environment, but that this effect is not exacerbated in the presence of Wnt-3a and therefore unlikely to underpin the changes in TNF-α release.

To further assess if the canonical β-catenin pathway is not involved in the exacerbation of TNF-α production, the canonical pathway inhibitor dickkopf-related protein 1 (DKK1) was added in culture for 24h, alongside all previous treatment groups. In cells that received DMSO vehicle, we observed the same patterns of TNF-α release as before (Figure 2C), namely a significant increase following stimulation with LPS (p<0.001), which was further exacerbated in cells treated with LPS + Wnt-3a (p<0.001). Importantly, no significant effects were found when the cultures were pre-treated with DKK1 compared to DMSO vehicle, with similar levels of TNF-α observed within each of the 4 groups (Figure 2C). We confirmed this was not due to an inability of DKK1 to reduce stabilisation of β-catenin, as addition of DKK1 on cells stimulated with Wnt-3a significantly lowered expression of β-catenin(Ser/33/37/Thr41) compared to Wnt-3a treated cells without DKK1 (Supplemental Figure 1), suggesting DKK1 was performing its expected function of inhibiting the canonical pathway.

### Wnt-3a mediated exacerbation of inflammation is dependent on the non-canonical pathways

The prior results highlight that the effects of Wnt-3a on inflammation may be independent of the canonical pathway. As inflammation has previously been linked to activation of the Wnt non-canonical pathways[1, 2], we therefore examined if the increased production of TNF-α following addition of Wnt-3a in an inflammatory environment was due to the involvement of these non-canonical pathways. To achieve this, inhibitors of each pathway were added to the culture 30 minutes before LPS/Wnt-3a.

As shown in Figure 3A, while LPS + Wnt-3a continued to result in greater levels of TNF-α compared to LPS alone (p<0.001) in vehicle treated cells, blocking the PCP non-canonical pathway with the JNK inhibitor SP600125 significantly reduced the increase in TNF-α triggered by LPS (p<0.01) and LPS + Wnt-3a (p<0.01). Similarly, blocking the calcium dependant pathway with U73122, a PLC inhibitor, not only decreased LPS-induced inflammation (Figure 3B p<0.01) but significantly decreased TNF-α levels in LPS + Wnt-3a treated cells (p<0.001). Together, these results suggest that the exacerbation of TNF-α production in an inflammatory environment is linked to non-canonical pathway activation rather than the canonical pathway.

**Figure 3:**
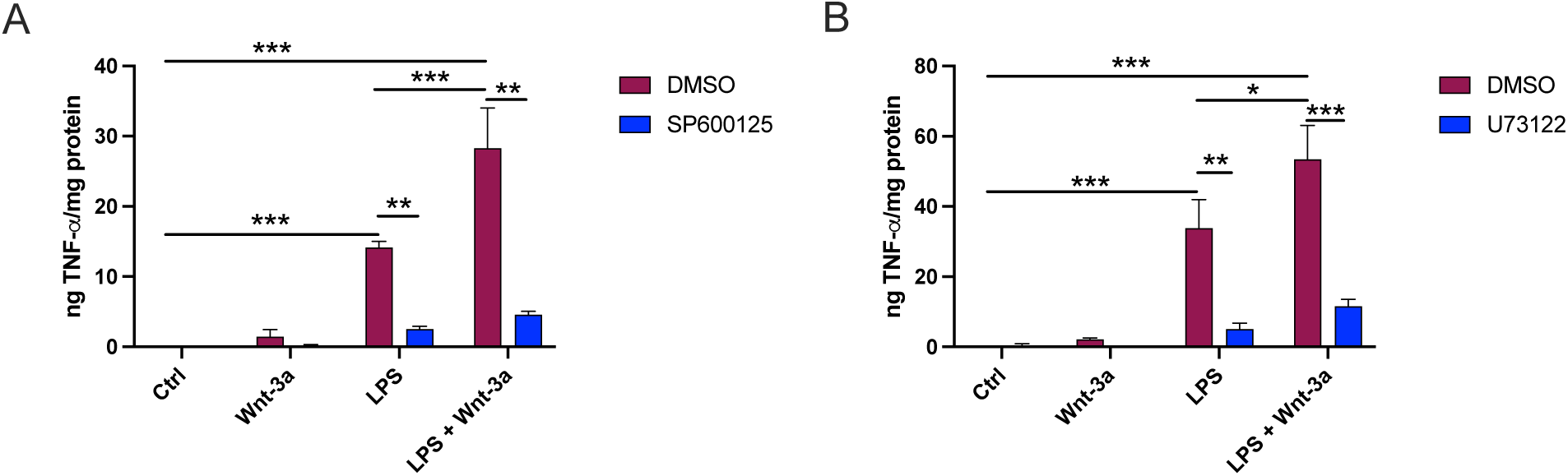
Inhibition of the two non-canonical pathways reduces TNF-α levels. Analysis of TNFα levels in microglia stimulated with the (A) JNK inhibitor SP600125 or (B) PLC inhibitor U73122. All values are shown as mean ± standard error of the mean (SEM). *p<0.05, **p<0.01, ***p<0.001. N=5/group.

### Wnt-3a does not alter DA neuron populations in the SNpc of MPTP lesioned mice

Our in vitro studies have highlighted the pro-inflammatory role of Wnt-3a in an inflammatory context. Since microglia-mediated neuroinflammation has been significantly linked to PD pathology[19–21], we examined if exogenous application of Wnt-3a would exacerbate dopaminergic neuron degeneration in the MPTP mouse model of PD, a model well established to induce inflammatory outcomes[12, 22, 23]. As anticipated, administration of MPTP led to a loss of TH positive neurons in the SNpc in vehicle (BSA) treated animals compared to saline controls (p<0.001). While Wnt-3a did not affect the baseline level of dopaminergic neurons, as indicated by no significant difference within saline-treated animals, it also failed to alter the number of surviving dopaminergic neurons against MPTP toxicity (Figure 4A, C). To ensure the accuracy of cell population estimates were not due to changes in TH expression, the number of NeuN positive cells was also quantified. Similarly, a significant reduction in NeuN-positive cells was observed after MPTP treatment (p<0.001) and the addition of Wnt-3a did not mitigate this effect (Figure 4B, D).

**Figure 4:**
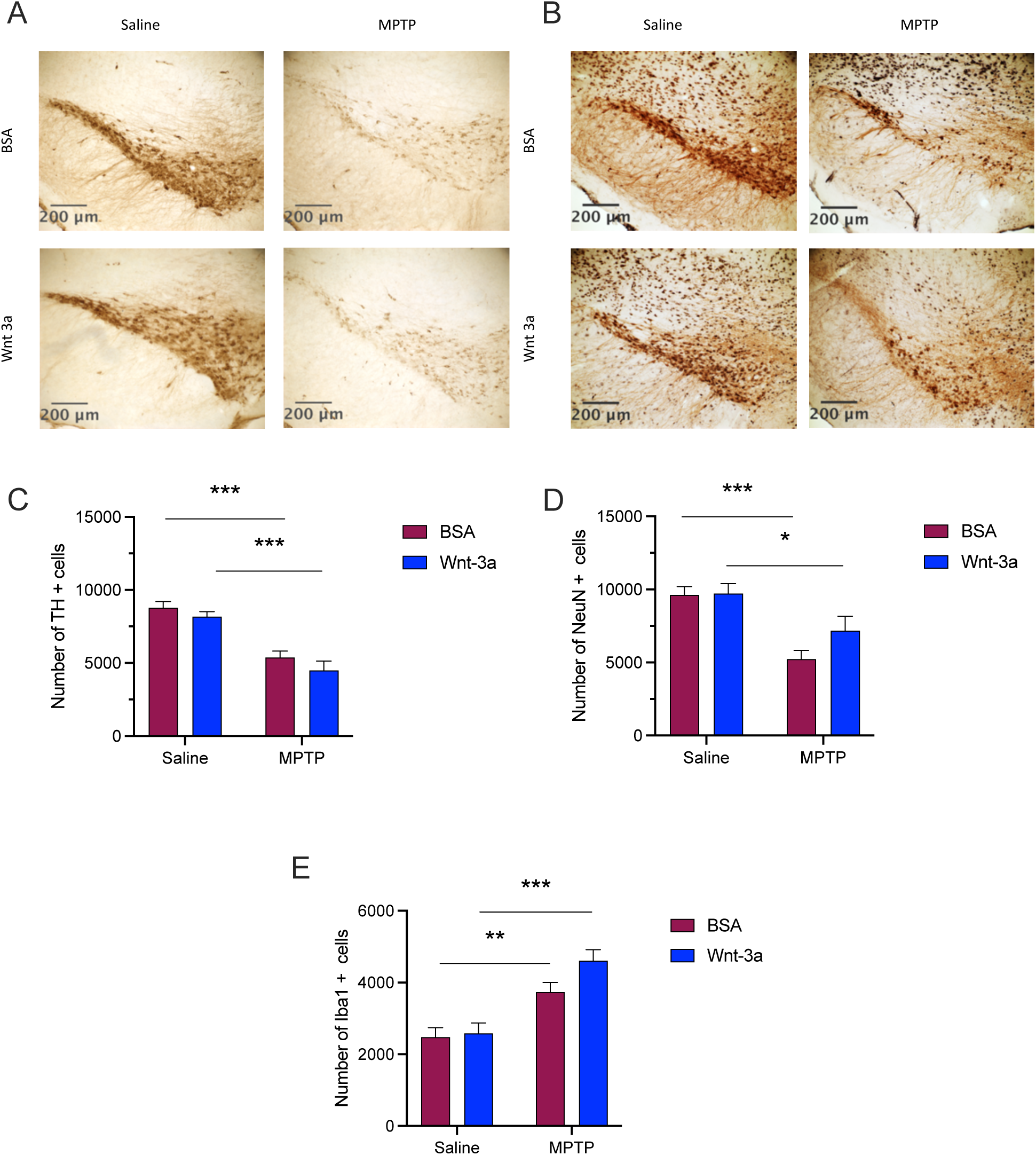
Wnt-3a in the MPTP mouse model of PD. Representative images of the substantia nigra pars compacta (SNpc) of (A) dopamine neurons identified by TH^+^ cells and (B) total neurons identified by NeuN^+^ cells, with TH used to outline SNpc region. Stereological quantification of (C) TH, (D) NeuN, and (E) Iba1 positive cells in the SNpc. All values are shown as mean ± standard error of the mean (SEM). *p<0.05, **p<0.01, ***p<0.001 compared to control. N= 7/group.

To determine if the inability of Wnt-3a to alter dopaminergic cell populations was due to any effects on microglia populations, we also quantified the number of Iba1+ cells in the SNpc via stereology. As expected, MPTP significantly increased the number of microglia in the SNpc (Figure 4E p<0.01) in vehicle treated animals. Although there were no changes in the baseline numbers of microglia as a result of Wnt-3a, animals that received MPTP + Wnt-3a had significantly higher numbers of Iba1 (p<0.001) positive cells compared to their saline controls. Importantly, there was no significant differences in microglia numbers between vehicle and Wnt-3a treated animals following MPTP, suggesting that exogenous application of Wnt-3a was unable to alter MPTP-induced inflammation.

## Discussion

Inflammation has been recognised as a significant contributor to neurodegenerative diseases, with microglia increasingly demonstrated to play a primary role in this process, making modulation of the inflammatory response an attractive target to mitigate disease-associated pathology[24–26]. Among the various strategies able to regulate inflammation, Wnts have gained increasing interest, particularly in the study of PD. Dysregulation of the Wnt canonical pathway in rodent models of the disease and in the brains of PD patients has been revealed, suggesting promoting an anti-inflammatory environment through the Wnt-mediated canonical pathway could be effective in decreasing neurodegeneration[27–31]. Although the precise effects of Wnts on inflammation are not yet fully understood, they appear to be influenced by the specific Wnt ligand, receptor, cell type and cellular environment. However, it is commonly assumed that stimulating the canonical pathway triggers an anti-inflammatory response, with studies showing this pathway can negatively regulate NFκB activity, reduce TNF-α release and induce the expression of the anti-inflammatory cytokine IL-10[32–34]. In contrast, the two non-canonical pathways have been shown to trigger inflammation, such as in human lung fibroblast, where Wnt-5b and Wnt-5a activate the PCP pathway, resulting in increased secretion of IL-6 and CXCL8 secretion along with NFκB activation[35]. Therefore, this study aimed to assess the effect of Wnt-3a, a classical canonical pathway activator, on primary microglia cultures. Interestingly, results suggest a pro-inflammatory glial response to Wnt-3a in an inflammatory context with little or no effect on non-stimulated cells.

To investigate the inflammatory response in glial cells, this study assessed the level of TNF-α, a pro-inflammatory cytokine linked to numerous neurodegenerative diseases, including PD [36, 37], following stimulation with Wnt-3a. Here, TNF-α production was exacerbated when Wnt-3a was added to reactive glial cells but not in cells displaying a resting state. Interestingly, not only is the NFκB pathway known to trigger pro-inflammatory cytokine secretion, but studies have shown an interconnection between the Wnt canonical pathway and NFκB, with β-catenin inhibiting NFκB while GSK3-β activates this pro-inflammatory protein[38]. We therefore examined key molecules associated with this pathway and notably, found no differences in phospho-IKKα/β, phospho-IκBα, NFκB, and NFκB(Ser536) between LPS and LPS + Wnt-3a groups, suggesting that the exacerbation is independent of NFκB pathway activation. Nevertheless, it is possible that changes in the NFκB pathway leading to increased TNF-α production occurred prior to the 24-hour post-stimulation measured in this study. Additionally, the contributions of other inflammatory pathways that intersect or interact with the Wnt pathway could also be considered. The map kinase (MAPK) pathway, for example, has been suggested to be linked to inflammation triggered by Wnt[39], with Wnt-3a stimulation of F9 and HEK293 cells leading to an increase of MAPK and P38 activity[40]. It is worth noting that a small increase of NFκB(Ser536) was observed in microglia stimulated with Wnt-3a alone. Previous studies have indicated, when not in a chronic state, neuroinflammation may be beneficial with low levels linked to immune surveillance, immune cell communication, development, memory, learning and remodelling[41]. Therefore, the small amount of active NFκB(Ser536) observed in this study could be an endogenous response of microglia to Wnt-3a stimulation that may serve as a beneficial function for neurons survival, however further studies would be necessary to determine the exact consequences of small amount of NFκB produced by immune cells on neurons.

It has previously been shown that activation and translocation of β-catenin to the nucleus has a role in decreasing inflammation[42]. It was expected that, as a canonical pathway activator, Wnt-3a would therefore decrease the LPS-induced released of TNF-α via its action on β-catenin. However, our study showed no difference in β-catenin(Ser33/37/Thr41), representing active β-catenin, between LPS and LPS + Wnt-3a groups. Therefore, the exacerbation of TNF-α appears to be independent of the canonical pathway. Indeed, when cells were exposed to the canonical inhibitor DKK1, the LPS + Wnt-3a group maintained an exacerbation of TNF-α production compared to the LPS group, confirming this effect was not due to dysregulation of the Wnt canonical pathway.

In contrast, the non-canonical pathways have been shown to participate in TNF-α production, with a previous *in vitro* study showing levels of activated JNK, a molecule found in the PCP pathway, were higher after LPS stimulated microglia compared to controls[43]. Additionally, Fyn, a kinase phosphorylating PKC (a member of the Ca^2+^ non-canonical pathway) has been shown to be required for TNF-α production by primary microglia stimulated with LPS[44]. Thus, inhibiting these pathways would be expected to lead to a decrease in inflammation. Indeed, the JNK inhibitor SP600125 has been shown to lead to a reduction in NO production and MAPK pathway in microglia stimulated with LPS[45], while U73122 can reduce pro-inflammatory cytokines production when added to cell lines and primary macrophages also stimulated with LPS[46]. Therefore, both non-canonical pathways were studied here for their role in the Wnt-3a-mediated exacerbation of TNF-α production. Inhibiting the Ca^2+^ pathway and the PCP pathway with U73122 and SP600125, respectively, resulted in a decrease in the Wnt-3a-mediated exacerbation of TNF-α. These results suggest that in reactive microglia, Wnt-3a stimulation results in a shift from activation from the canonical to non-canonical pathways, exacerbating the previously identified role of the non-canonical pathway in inflammation following LPS stimulation. As Wnt5a, which is classically associated with activating the non-canonical pathways, is able to dose-dependently increase TNF-α mRNA levels in macrophages[47], it would be of interest to determine if exogenous application of Wnt-3a in an inflammatory context as performed in our study leads to an increase in Wnt5a levels, which may underscore the exacerbation of TNF-α release found here.

Activating the canonical pathway in immune cells as a mechanism to promote an anti-inflammatory response has been shown to protect dopaminergic neurons against cellular death. In a previous study utilising the MPTP model of PD, this neurotoxin was found to reduce the activity of the canonical pathway in astrocytes, preventing the ability of dopamine neurons to recover after injury while activating the canonical pathway using a GSK3β inhibitor was neuroprotective[9]. Moreover, blocking the canonical pathway by unilateral infusion of a β-catenin antagonist in the SN leads to reactive astrocytosis and inhibits TH^+^ neuron survival in a MPTP-mouse model[17]. In a 6-OHDA rat model, Dun et al., showed that treatment with DKK1 led to aggravation of dopaminergic neuronal damage in the SNpc while pre-treatment with a GSK3β inhibitor for 7 days resulted in neuroprotection[16]. More recently, changes in endogenous Wnt signalling were found in LRRK2 knockout and LRRK2 G2019S knockin mouse lines compared to controls[48]. Together these results suggest a role of the canonical pathway in degeneration in PD. Interestingly, in our in vivo MPTP experiments, exogenous application of Wnt-3a directly into the brain was unable to protect against MPTP-induced dopaminergic cell death with a similar inability to alter numbers of microglia in the SNpc. While these results may be considered surprising based on the previous studies outlined above noting a dysregulation of the canonical pathway in PD, it should be noted these studies manipulated the canonical pathway activation via DKK1 addition or GSK3β inhibition, therefore the effects of the canonical pathway were studied but not the direct and more upstream effects of exogenous Wnt in the brain of a PD animal model.

This research has certain limitations that need to be considered. While the primary focus was on TNF-α, given its crucial role in neuroinflammation and neurodegeneration, it would be valuable to explore how other pro-inflammatory cytokines, such as IL-1β or IL-6, align with the observed TNF-α profile seen here after Wnt-3a stimulation. Additionally, examining the response of anti-inflammatory cytokines such as IL-4 or IL-10 following Wnt-3a stimulation in resting vs inflammatory cells could provide complementary insights. Second, this study reveals an exacerbation of inflammation in the LPS + Wnt-3a group after 24h. Typically, β-catenin levels are already elevated after 2-3 hours following Wnt-3a stimulation and remain stable at later timepoints[49]. To obtain a comprehensive understanding of the inflammatory response, it would be important to assess the outcomes of exogenous Wnt-3a on β-catenin and other upstream and downstream proteins associated with Wnt signalling at earlier timepoints. Several studies have demonstrated a decrease of neuronal viability following application of culture medium harvested from glial cells that were stimulated with LPS. Thus, a similar approach could be implemented to investigate the role of Wnt-3a on this relationship over time and if the exacerbation of inflammation in reactive glial cells leads to a decrease in neuronal viability. Finally, this study primarily focused on the effect of Wnt-3a in resting and activated microglia, given the crucial role of this immune cell in the pathophysiology of PD, however determining if a similar exacerbation of pro-inflammatory cytokine release occurs in activated astrocytes and if this effect is mediated by the non-canonical Wnt pathway would be of interest.

All together, these results show the importance of the cellular context in the Wnt response. Wnt-3a, a well-known activator of the canonical pathway, can shift and activate different pathways in an inflammatory environment, leading to an increase of pro-inflammatory cytokine production. Therefore, understanding the molecular mechanisms that are able to predict the outcomes of addition of Wnts is of high importance in the context of developing pertinent therapies for neurodegenerative diseases.

## Acknowledgement

The authors would like to thank members of the Centre for Neuroscience and Regenerative Medicine for their technical support and assistance in editing this manuscript.

## Author contributions

GF, SS, PR, and BV conceptualised the studies. GF and SS participated performed the studies. SS and BV provided supervision. GF, SS, PR, and BV wrote and edited the manuscript.

## Availability of data and materials

The datasets used and/or analysed are available from the corresponding author on reasonable request.

## Declarations

## Ethical approval and consent to participate

This study was conducted in accordance with theAustralian National Health and Medical Research Council animal experimentation guidelines and the local Code of Practice for the Care and Use of Animals for Scientific Purposes (2004). All experiments were approved by the Garvan Institute Animal Ethics Committee under approval numbers 18/16 and 17/35.

