## Supplemental Figure 1 for "Wnt-3a exacerbates production of TNF-α in LPS stimulated microglia independent of the β-catenin canonical pathway"

### Supplementary Information

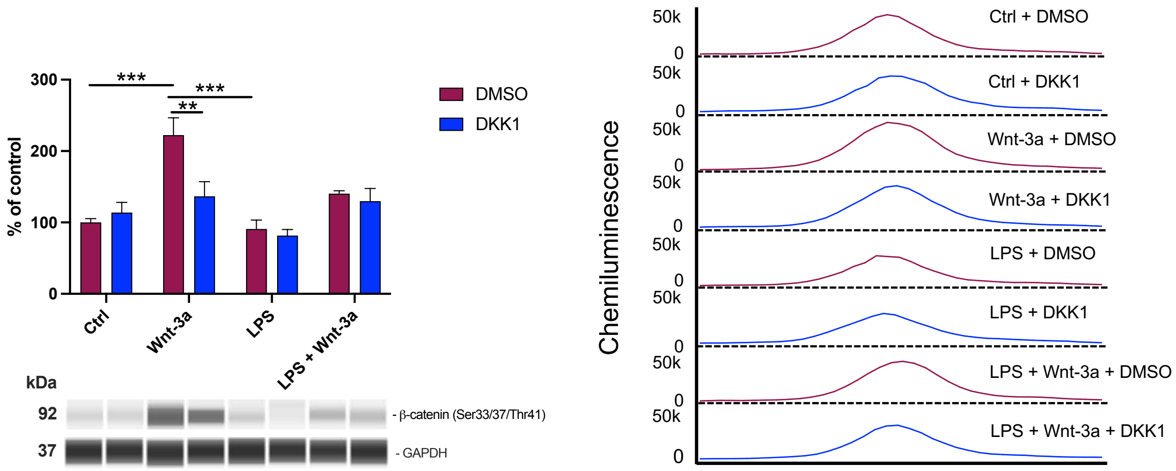


**Supplemental Figure 1.** Relative expression changes, corresponding Wes-generated pseudobands and Wes-generated pseudobands of active β-catenin in microglia stimulated with LPS and/or Wnt-3a and PBS or the canonical pathway inhibitor DKK1. All values are shown as mean ± standard error of the mean (SEM). **p<0.01, ***p<0.001. N=5–6/group
